# Gray matter abnormalities in sight deprivation and sight restoration

**DOI:** 10.1101/2024.11.29.626016

**Authors:** Caterina A. Pedersini, Alessio Fracasso, Amna Dogar, Bas Rokers, Pawan Sinha

**Affiliations:** Psychology, New York University Abu Dhabi, United Arab Emirates; School of Psychology and Neuroscience, University of Glasgow, Hillhead Street 62, Glasgow G12 8QE5, Scotland, UK; ASPIRE Precision Medicine Research Institute, Abu Dhabi, United Arab Emirates; Department of Brain and Cognitive Sciences, Massachusetts Institute of Technology, Cambridge, Massachusetts, United States of America

## Abstract

Blindness provides a unique model for investigating brain plasticity in response to sensory deprivation. While structural changes in both gray and white matter have been widely documented, particularly in cases of early or congenital visual deprivation, gray matter studies have traditionally focused on cortical thickness, often finding cortical thickening in posterior regions. However, other aspects of gray matter integrity, such as cortical myelin content, remain underexplored. In this study, we examined the effects of visual deprivation on cortical structure in a cohort of congenitally blind individuals who received eye surgery during adolescence, expanding beyond conventional measures to include cortical thickness, curvature, and T1-weighted signal intensity. This multi-faceted approach offers a more comprehensive view of cortical adaptations to congenital sensory deprivation.

While blindness offers valuable insights into sensory-driven brain plasticity, an intriguing and unresolved question is whether structural plasticity reverses after sight restoration, enabling typical visual processing circuits to develop despite the initial period of deprivation. To address this, we assessed the effect of sight-recovering eye surgery on gray matter changes. Critically, individuals in this cohort received surgery after the closure of the sensitive period for visual development. We did not find evidence of gray matter changes after surgery. However, in a previous study conducted on the same cohort, we reported that notable plasticity in white matter emerged in this same population. These results suggest that white matter alterations, rather than gray matter changes, may potentially serve as a biomarker of structural plasticity following sight restoration, even beyond the sensitive developmental window.

## Introduction

Blindness offers a unique opportunity to study functional and structural changes following sensory deprivation in the absence of brain lesions. Functional plasticity in blindness has primarily been examined through two lenses: (1) the functional specialization of brain regions during non-visual tasks and (2) the integration of unimodal sensory regions into broader brain networks via functional connectivity (FC) analysis. In the case of functional specialization, early blind individuals (EBs) exhibit either increased activity in non-deprived sensory areas (Elbert et al., 2002; Hertrich et al., 2013) or processing of non-visual sensory information within occipital regions traditionally associated with vision (Frasnelli et al., 2011; López-Bendito et al., 2022). In terms of network integration, numerous studies have reported changes in both stationary (Abboud & Cohen, 2019; Bauer et al., 2017; Burton et al., 2014; Hou et al., 2017; Hu et al., 2020; Huang et al., 2020; Liu et al., 2007; Striem-Amit et al., 2015; Wen et al., 2018) and dynamic FC (Pelland et al., 2017) following early sensory deprivation. Recently, we employed Hidden Markov Models (HMM) on resting-state fMRI (rs-fMRI) data from EBs, and showed that early visual deprivation alters FC temporal dynamics (Pedersini et al., 2024). Our findings revealed that EBs use to visit more than controls brain states that primarily activate unimodal regions, with heightened connectivity involving occipital areas, although the overall frequency of switching between brain states remains intact. Interestingly, the altered frequency of visiting specific states offers insights into stationary FC patterns following sensory deprivation, highlighting the role of the temporal dynamics in defining patterns of FC. These findings highlight the profound impact of visual deprivation during critical periods of visual development (Röder et al., 2021), reshaping functional brain organization and affecting both deprived and non-deprived unimodal sensory regions.

Structural plasticity in blindness has been examined by comparing alterations in gray and white matter following early sensory deprivation. Regarding white matter, several studies have documented altered structural integrity in retinofugal projections (Ankeeta et al., 2021; Ptito et al., 2021) in EBs. Gray matter alterations have traditionally been studied using either Voxel-Based Morphometry - VBM (Anurova et al., 2015; Jiang et al., 2009; Voss et al., 2014) or Surface Based Morphometry - SBM (Anurova et al., 2015; Bauer et al., 2017; Inuggi et al., 2020). These studies indicate that in EBs, gray matter tends to be thicker in posterior cortical areas compared to sighted controls, potentially due to reduced axonal pruning and synapse elimination during visual development (Jiang et al., 2009). A recent meta-analysis (Paré et al., 2023) further highlighted these structural differences at both whole-brain and ROI-level. Noted limitations of these previous studies are related to the technique used - VBM does not easily allow differentiation of group differences into specific cortical features such as thickness, curvature, or T1-weighted (T1-w) signal intensity - and to the metrics extracted - SBM studies have predominantly focused on cortical thickness and gray matter volume. To our knowledge, only one study (Voss et al., 2014) examined cortical myelin content through magnetization transfer imaging, reporting increased myelination in posterior areas when compared to controls. To address these limitations, the primary goal of this study is to explore the impact of congenital sensory deprivation on cortical structure in adolescents, extending beyond traditional findings by focusing on cortical thickness, curvature, and T1-weighted signal intensity. This approach acknowledges that, while myelinated axons are mainly found in white matter, some are also present in the cortex, contributing to cortical myeloarchitecture. The T1-w signal intensity can serve as a proxy for myelin within gray matter, as it reflects both myelin and iron, with the latter being highly co-localized with the former (Fukunaga et al., 2010). T1-w signal intensity has been used to map intracortical gray matter features in striate (Barbier et al., 2002; Clare & Bridge, 2005; Fracasso et al., 2016) and extra-striate cortex (Walters et al., 2003), and to study whole-brain cortical myelination patterns (Fracasso et al., 2016; Lutti et al., 2014; Sereno et al., 2013)

An intriguing and still unresolved question is whether brain plasticity reverses after sight restoration, allowing typical visual processing circuits to emerge despite an initial phase of congenital sensory deprivation (Bottari et al., 2016; Sinha & Held, 2012). In task-related brain activity, Bottari and colleagues (2018) observed that compensatory plasticity - such as enhanced auditory motion processing - appears to persist even after sight restoration, while visual motion perception remains impaired. Other studies have confirmed that auditory-driven activation of early visual areas persisted after sight-recovery surgery and tended to decrease over time following the procedure (Collignon et al., 2015; Dormal et al., 2015). In resting-state fMRI, Rączy and colleagues (2022) suggested a partial retraction of cross-modal plasticity following sight restoration, indicating an incomplete recovery of typical visual resting-state activity patterns, which may form the basis for persistent impairments even after sight is restored. These findings support the notion that the excitatory/inhibitory (E/I) balance, which has shown to be disrupted following congenital sensory deprivation (Muret & Makin, 2021), may remain altered even years after sight-restoration surgery (Pant et al., 2024).

Structural changes following sight restoration remain less explored. In 2015, Guerreiro and colleagues reported increased cortical thickness in occipital areas, which negatively correlated with behavioral performance on an audio-visual task, despite the transient nature of the early sensory deprivation. However, in this study, the clinical cohort was composed of individuals who received the eye surgery before the 2nd year from birth, thus before the closure of the sensitive period of visual development, raising an intriguing question: how does structural brain plasticity reverse after sight restoration received during adolescence? In a recent study, we found that sight recovery following eye surgery received during adolescence induced structural plasticity in late-visual white matter pathways (Pedersini et al., 2023), underscoring that white matter tracts are sensitive markers of structural brain plasticity following vision restoration. To further investigate anatomical changes following eye surgery, in this study we assessed gray matter alterations in the same clinical cohort of congenitally blind individuals who underwent eye surgery after the closure of the sensitive period of visual development.

## Methods

### Participants

Data acquisition was performed according to the Code of Ethics of the World Medical Association (Declaration of Helsinki, 2008) and was approved by the Institutional Review Boards of Dr. Shroff’s Charity Eye Hospital (Delhi, India) and the Massachusetts Institute of Technology (Cambridge, Massachusetts, United States). Informed consent was obtained from all individuals who have been treated for dense bilateral congenital cataract (congenital cataract-reversal individuals - CC). All participants were evaluated and treated by licensed medical professionals at Dr. Shroff’s Charity Hospital in Delhi, India. Twenty-three individuals with cataracts (6 female), aged 7 to 21 years (12.34 ± 3.69 years at the time of surgery) were enrolled in this study. They were screened and recruited from Uttar Pradesh, India, through Project Prakash, which provides ophthalmic care to children in rural areas. All participants had dense, bilateral cataracts present since before one year of age. Prior to receiving surgery to remove the natural lens and implant an artificial intraocular lens (IOL) in each eye, they had not received any eye care. Following surgery, additional vision correction was provided through glasses and other low-vision aids. Participants’ ages were reported to the nearest year by their parents or guardians. Due to limited record-keeping in rural families, exact birth dates were unavailable, and all analyses were based on the closest estimated age at the time of surgery. More details about participants’ selection and recruitment can be found in (Pedersini et al., 2023). Structural images were acquired at multiple timepoints for the majority of CC individuals (15) for a total of 58 sessions across 23 individuals. Two sessions were excluded from subsequent analysis due to problems with brain extraction during data preprocessing. Moreover, the first session of subject P06 was removed from the analysis due to the low quality of the structural image, as reported by MRIQC (Esteban et al., 2017) (see fig. S1). Additionally, 172 healthy normally sighted controls (NSC) were included by selecting 10 age- and gender-matched controls for each patient and session from the 3T Human Connectome Project in Development dataset (HCP-D, https://www.humanconnectome.org/study/hcp-lifespan-development) (Harms et al., 2018; Somerville et al., 2018) (mean age of 13.67 ± 4.45 years at the time of data collection).

### Magnetic Resonance Imaging Data Acquisition

Brain imaging data of patients were obtained at the Mahajan Imaging Center, Defense Colony (New Delhi, India) using a GE Discovery MR750w 3T MRI scanner (GE Healthcare, Inc, Chicago, IL, USA) equipped with a 32-channel head coil. For each participant, either one or multiple structural whole-brain T1-weighted anatomical scans (3.7 ms TE; 9.5 ms TR; 1×1×1 mm^3^ isotropic voxels) were acquired. Details of the HCP-D brain imaging acquisition protocol are extensively described in (Harms et al., 2018).

### Data Processing and Analysis

We used custom algorithms to optimize skull-stripping and minimize the need for manual correction. For each sight-recovery individual and session, we performed ANTs as well as BET (FSL) with 3 different fractional intensity thresholds (0.2, 0.3 and 0.4), and selected the best result based on visual inspection. The remaining steps of recon-all were then performed on these skull-stripped images (*-autorecon1 -noskullstrip*) to segment the surface of each subject in native and fsaverage space. Manual cleaning was performed to correct for possible errors in the surface segmentation, either by refining the brain mask or by adding control points. AD and CAP (both authors of this study) were in charge of the manual cleaning, with AD performing it and CAP supervising the process. Data from the HCP-D were included in the study without performing any additional manual cleaning.

We extracted cortical thickness (CT), curvature (CV), and T1-weighted (T1-w) signal intensity, used as a proxy for cortical myelin content (Fracasso et al., 2016; Glasser & Van Essen, 2011), at each vertex of the whole brain, to assess group differences. T1-w signal intensity was obtained at the mid-space surface, which lies between the pial and white matter surfaces, using the *mri_vol2surf* function that generates a vector of T1-w signal values for each vertex. CT and CV were automatically extracted using the recon-all function. All structural metrics were computed in fsaverage space without applying any smoothing.

T1-weighted signal intensity can vary depending on the scanner’s scaling factor, and is not a quantitative measure. Since data from sight-recovery and sighted participants were collected at different sites using different scanners, it was essential to control for scanner-related variability. When plotting mean T1-weighted signal intensity, cortical thickness (CT), and curvature (CV) across participants, we observed a global group effect (Fig. 1A), with T1-w signal and CT being generally lower in the sight-recovery group compared to sighted participants. These differences could be attributed to scanner variability, manual cleaning procedures, differences in software versions or basic differences between groups, such as education and socio-economic status. To mitigate the impact of these factors and to control for global differences between subjects, we standardized each structural measure for each participant and hemisphere, using the formula:

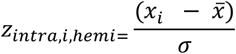

where *x*_*i*_ is the structural metric for the *i*-th vertex in a hemisphere, 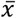 is the average across vertices in that hemisphere, and σ is the standard deviation across vertices of the same hemisphere. The same procedure was applied for both hemispheres.

**Fig. 1.**
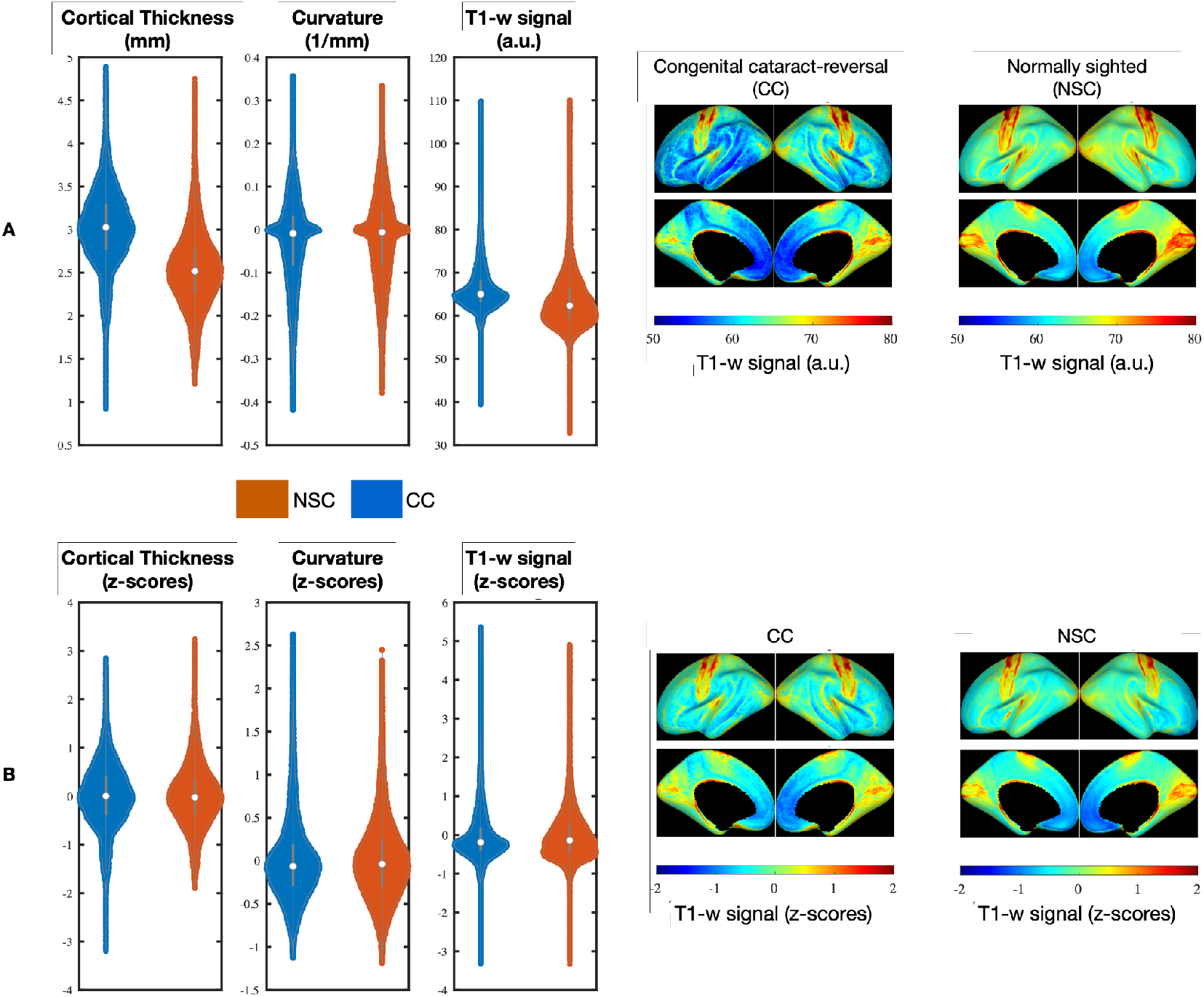
Distribution of cortical thickness (CT), curvature (CV) and T1-w signal intensity across participants. A - Left: Violin plot showing CT, CV and T1-w signal for normally sighted controls (NSC) and congenital cataract reversal (CC) individuals, using the raw data. A - Right: lateral and medial brain surface representing the mean T1-w signal intensity for CC and NSC, using the raw data. B - Left: Violin plot showing standardized values of CT, CV and T1-w signal for CC and NSC. B - Right: lateral and medial brain surface representing the mean standardized T1-w signal intensity for CC and NSC.

T1-weighted signal intensity is influenced by cortical thickness, due to partial volume effects, tissue composition variations, as well as by curvature (Annese et al., 2004; Sereno et al., 2013). To account for these factors, we regressed out the contributions of cortical thickness (CT) and curvature (CV) from the T1-w signal intensity of each subject and session, by fitting the following linear model:

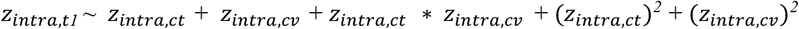

where *z*_*intra*,*t1*_ represents the z-scored T1-w signal, *z*_*intra*,*ct*_ is the z-scored cortical thickness, and *z*_*intra*,*cv*_ is the z-scored curvature. The residuals and intercept were retained for subsequent analysis as a proxy for myelin content. The following steps involve two paths: 1) assessing structural alterations resulting from early blindness, to evaluate structural brain plasticity following a congenital visual impairment and 2) evaluating the impact of sight-recovery surgery on brain structure, to assess how the eye surgery may affect those structural alterations.

#### Assessment of structural alterations following congenital blindness

We performed a whole-brain mass-univariate linear mixed-effects model to investigate structural changes in cortical thickness, curvature, and T1-weighted signal (a proxy for cortical myelin content) in CC participants, following the congenital sensory deprivation. For each vertex, we fitted a linear mixed-effects model that included one fixed-effect of interest (group), two fixed-effects of no interest (age at measurement and gender), and a random intercept for subjects to account for multiple timepoints per participant:

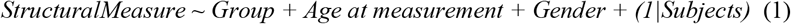

For each vertex, we evaluated the fixed effect of interest (Group). We applied a cluster correction using the function *mri_glmfit-sim*, with a vertex-wise cluster threshold of 3 (p = 0.001) and a cluster-wise p-threshold of 0.05. Additionally, we performed the same linear mixed-effects model at the ROI level using the Desikan-Killiany atlas (Desikan et al., 2006). For each subject and region, we extracted the median z-score for each structural measure, resulting in a 227 (acquisitions) by 34 (ROIs) matrix. These median values were entered into the linear mixed-effects model to assess between-group differences at the ROI level. P-values were corrected for multiple comparisons using FDR (alpha = 0.01), separately for each structural measure. It is important to note that we used standardized structural values as dependent variables, which means we assessed differences in the deviation of each measure from the within-subject mean. We visualized the whole-brain beta values resulting from the models on a brain surface, where warm (cold) colors indicate higher (lower) values in sight-recovered participants. These visualizations were generated after applying three different levels of smoothing: 0, 5, and 10 mm FWHM. The analysis at the ROI level was performed without smoothing.

#### Assessment the impact of sight-recovering surgery on brain structure

The primary goal of this analysis was to determine whether sight-recovery surgery could lead to structural brain changes, even when performed during adulthood. To test this, we performed a linear mixed-effects model on the group of sight-recovery individuals, at both the whole-brain and ROI levels. The model we applied was:

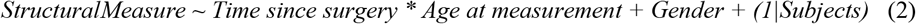

where all timepoints at or prior to the day of surgery were set to 1 to ensure a log10 value of 0, and the log10 of the number of days since surgery was calculated for all subsequent timepoints. The dependent variable was either the set of vertex-level values (for whole-brain analysis) or the median value across all vertices in the same ROI, merging data from the left and right hemispheres. For this analysis, p-values were corrected for multiple comparisons using FDR (alpha = 0.01), separately for each structural measure. We used the same model in a previous study (Pedersini et al., 2023), where we focused on assessing structural brain plasticity in terms of white matter integrity following the sight-recovering eye surgery. We fitted Bayesian linear models and obtained Bayes factors using the same model specification as in Equation (2). This approach allowed us to compare the influence of time since surgery with a null model that included only the remaining covariates. In describing our results, we adopt the terminology previously introduced regarding evidence in favor of or against the null hypothesis (Lee & Wagenmakers, 2014; Jeffrey, 1939). Specifically, we categorize the evidence as “weakly in favor/against” (Bayes factors <3), “moderately in favor/against” (Bayes factors 3–10), and “strongly in favor/against” (Bayes factors >10).

## Results

### Assessment of structural alterations following congenital blindness

The primary goal of this study is to assess structural changes associated with congenital blindness, focusing on cortical thickness, curvature, and T1-w signal intensity as a proxy for cortical myelin content. We observed large-scale structural changes following the congenital sensory deprivation, predominantly in cortical thickness and T1-w signal intensity.

Regarding the former, we found that CC individuals exhibited a thicker cortex in posterior regions following the congenital visual deprivation, along with scattered differences across the whole brain (see Fig. 2), with generally thicker CT along the medial and thinner CT on the lateral brain surface. It is important to consider that, as we are using standardized values, our results express departures from the individual subject mean, regardless of absolute differences between groups. These findings align with previous studies, which have reported gray matter structural changes following congenital visual deprivation, mainly in posterior regions. The analysis at ROI-level (see Fig. 3) pointed to the lingual gyrus and the anterior cingulate cortex as the two main regions showing thicker CT in CC than NSC, confirming previous findings (Anurova et al., 2015; Inuggi et al., 2020). Interestingly, we observed thicker CT in NSC in the temporal pole, a complex anatomical region associated with multiple high-level cognitive and visual functions (Tanaka, 1996).

**Fig. 2.**
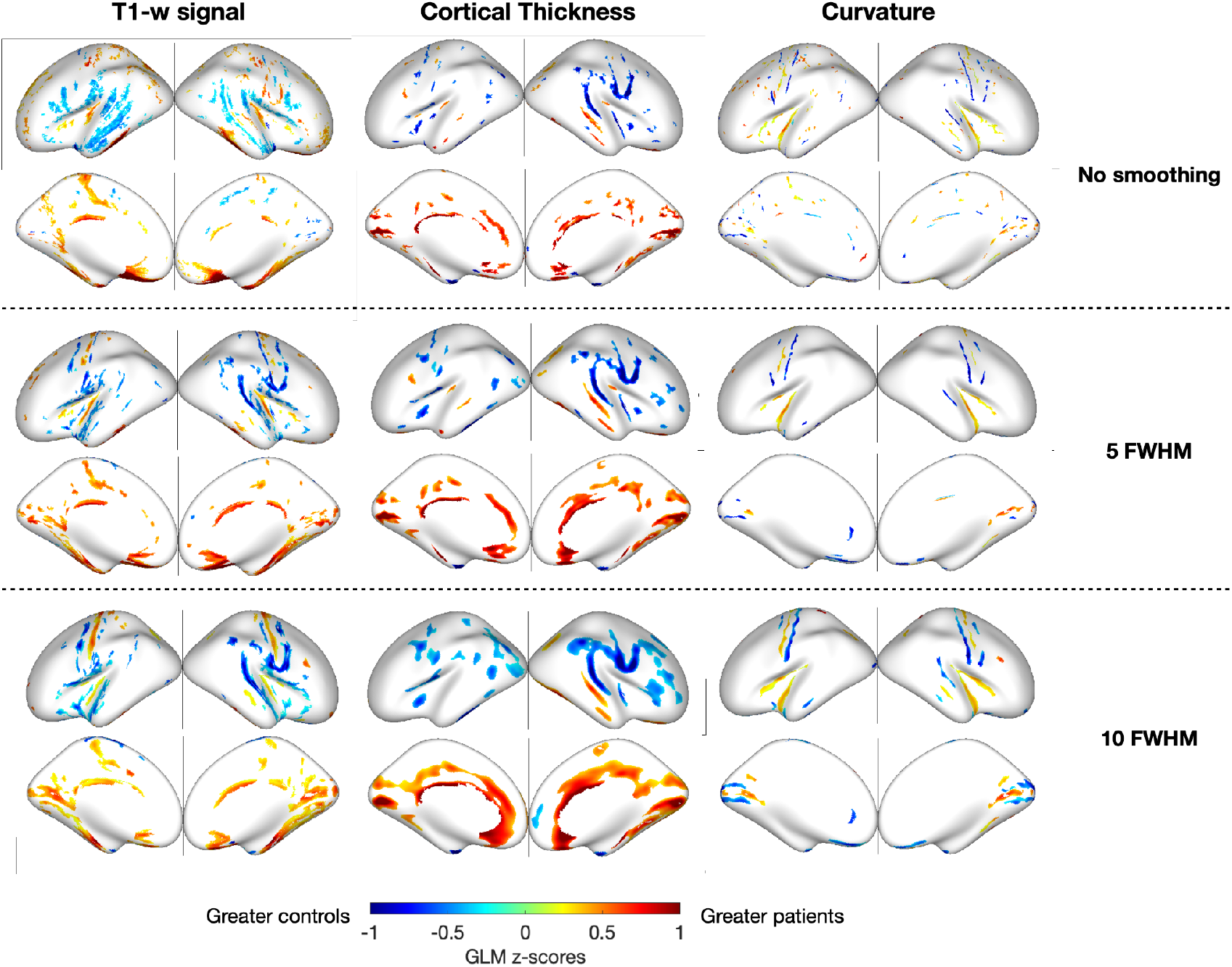
Effect of group at whole brain level. Cold colors represent z-scores greater in normally sighted controls (NSC); warm colors represent z-scores greater in congenital cataract-reversal (CC) participants. Results are shown after applying different levels of smoothing. Results are significant after cluster correction (vertex-wise cluster threshold of 3 and a cluster-wise p-threshold set to 0.05). CC individuals show greater T1-w signal and thicker thickness in occipital and callosal cortex, compared to NSC.

**Fig. 3.**
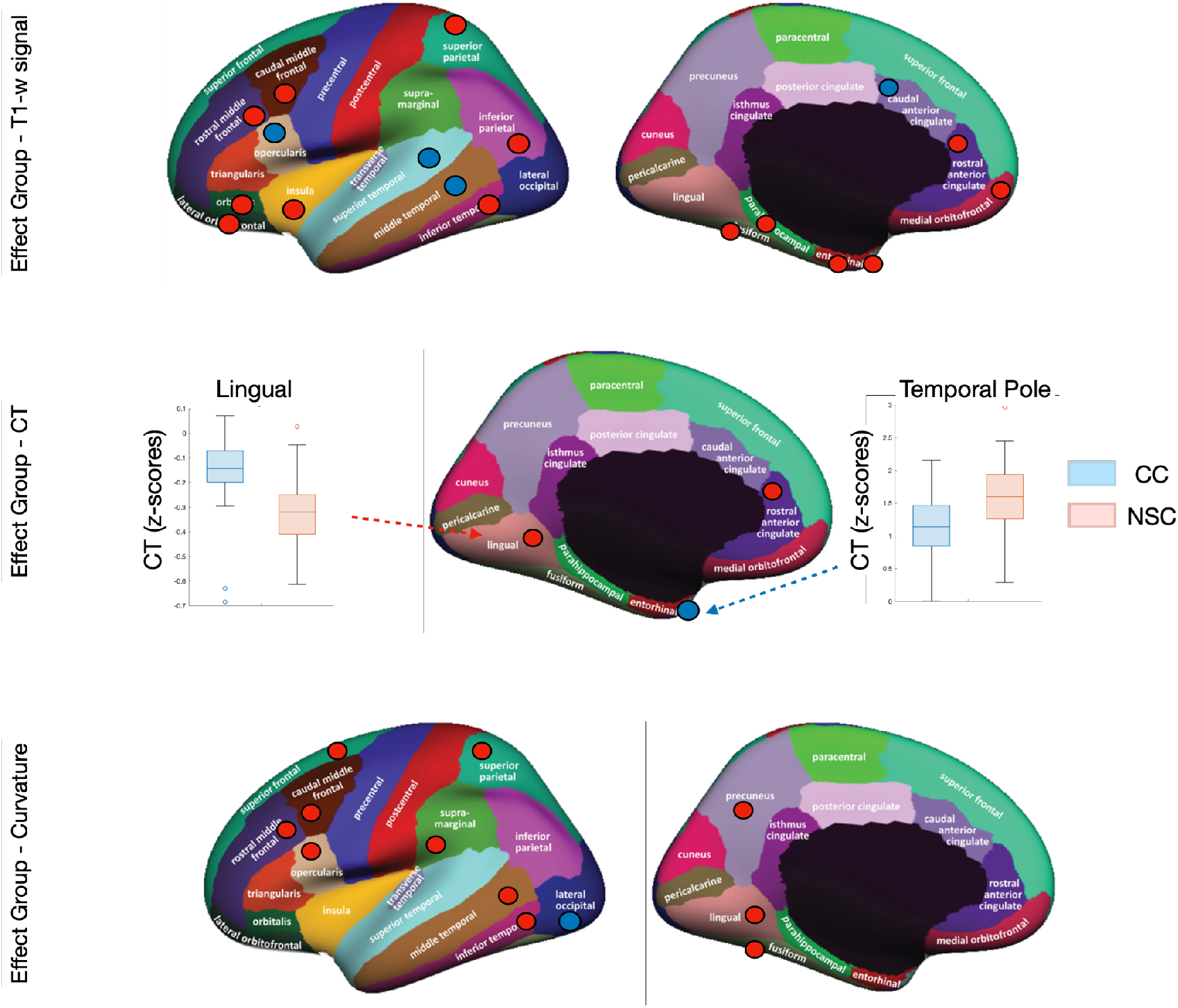
Effect of group at ROI-level on a brain surface. Visualization of the Desikan-Killiany ROIs on medial and lateral brain surfaces. Red dots highlight regions with significantly greater values in CC individuals, whereas blue dots indicate regions with significantly greater values in NSC. The boxplot in the central panel indicates z-scores cortical thickness for CC (blue) and NSC (red) participants in two main ROIs, in which we observed a significant effect of the group.

Regarding the latter, CC exhibited significantly greater T1-w signal intensities in the lateral occipital, parietal, and frontal regions compared to controls, relative to their individual means (see Fig. 3 and S2). Notably, we observed an increased proxy for myelin content in extrastriate brain regions associated with both the ventral and dorsal visual pathways, as well as in frontal regions, particularly middle and orbitofrontal areas. These findings underscore a more widespread pattern of structural changes compared to the results of Voss et al. (2014), reporting more localized between-group differences in magnetization transfer imaging, primarily within posterior regions.

When examining curvature, we found significant differences at ROI-level, with greater deviations from each subject’s average brain surface curvature in CC than NSC, particularly in occipital, temporal, and parietal regions.

A similar pattern was observed when considering only timepoints either within 10 days following sight-restoring surgery or well after the surgery (more than 10 days post-operation) (see Fig. S3), suggesting that the between-group differences don’t reverse after the sight-recovering eye surgery. Finally, we plotted the trend of between group differences across the range of ages and observed that the differences between NSC and CC are stable in time (see Fig. S4).

### Assessment of the impact of sight-recovered surgery on brain structure

The second goal of this study is to assess structural changes following the sight-recovering eye surgery, focusing on cortical thickness, curvature, and T1-w signal intensity as a proxy for cortical myelin content, while controlling for the effect of age (see Fig. S5). Interestingly, we did not observe any significant structural changes or reductions in gray matter after the surgery, either at whole-brain or ROI-level.

Fig. 4 shows the results from the ROIs reporting the lowest p-values for each structural measure. The BIC-lme Bayes Factor (BF) favors the null model, suggesting that the inclusion of time since surgery as a fixed effect does not significantly improve the model’s fit. This indicates that time since surgery does not play a significant role in explaining the variability of the dependent variables. Most BF values are below 0.33, indicating a preference for the null model over the alternative model (which includes time since surgery as a fixed effect). In some regions, the BF falls between 0.33 and 3, suggesting only anecdotal evidence in favor of one model over the other. These regions include 1) T1-w signal in the entorhinal cortex (BF = 0.34), 2) cortical thickness in the posterior cingulate (BF = 0.42), and 3) curvature in the frontal pole (BF = 0.38), inferior temporal (BF = 0.6), lateral occipital (BF = 0.44), lateral orbitofrontal (BF = 2.04), pars orbitalis (BF = 2.10) and rostral anterior cingulate (BF = 0.35). Only the caudal middle frontal region reported a Bayes Factor above 3 for curvature (BF = 6.3), indicating moderate evidence in favor of an effect of time since surgery (uncorrected p-value = 0.0016; FDR-corrected p-value = 0.0542).

**Fig. 4.**
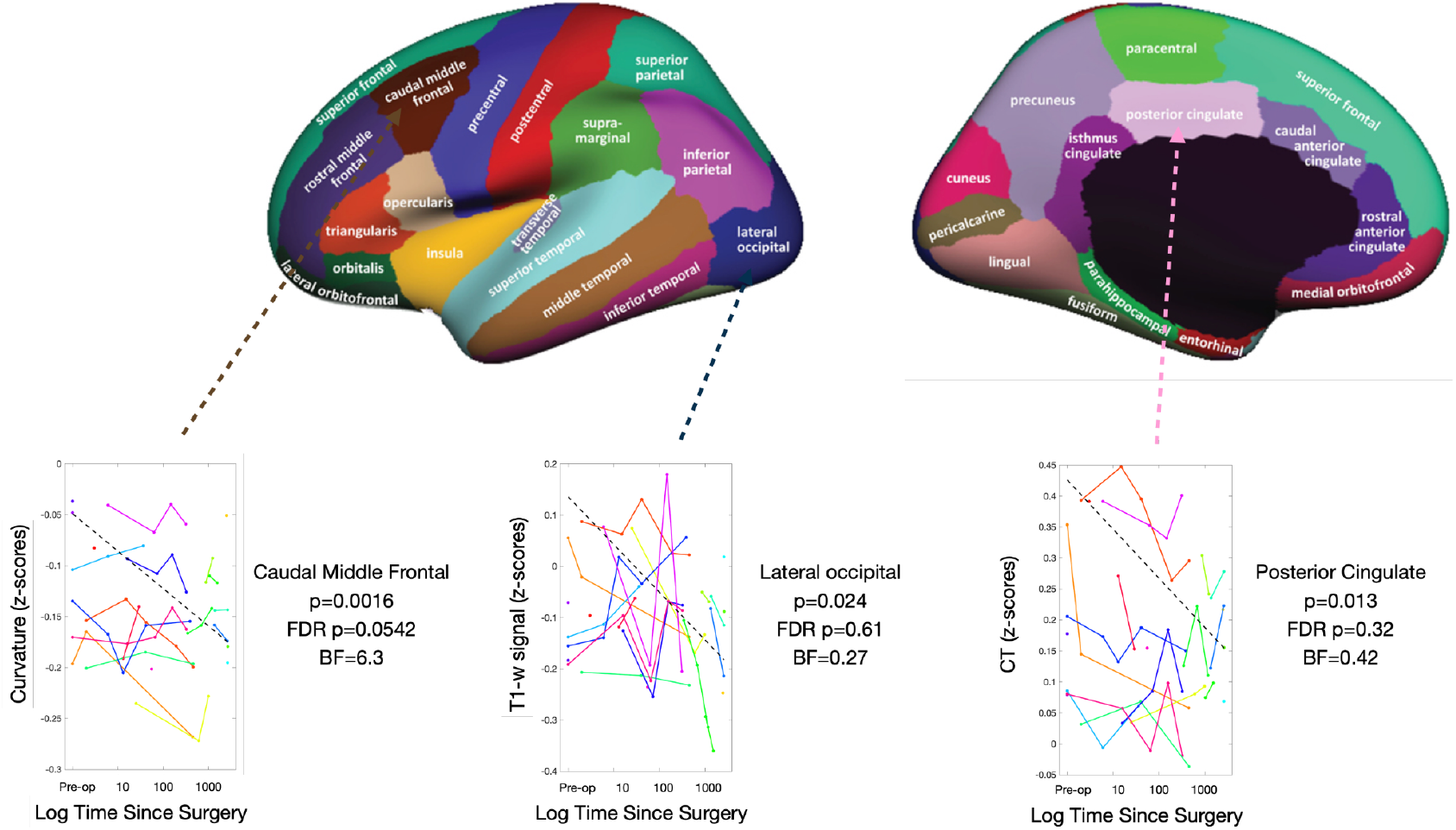
Effect of time since surgery at ROI-level in the group of individuals who have been treated for dense bilateral congenital cataract. Lateral and medial brain surface superimposed with the Desikan-Killiany Atlas. Each plot represents z-scores T1-w signal, cortical thickness or curvature for each subject as a function of time since surgery. Each color represents a CC individual and each line is connecting the multiple timepoints from the same subject. The dashed line indicates the beta extracted from the linear mixed-effect model, with the random-intercept of subjects. The plots have been chosen based on the ROIs reporting the lowest uncorrected p-value for the effect of time since surgery and the highest Bayes Factor (caudal middle frontal for curvature). None of the p-values survived the FDR correction.

In summary, these findings suggest no evidence of structural brain plasticity in gray matter following the sight-recovering eye surgery. It is essential to reconcile these results with the improved behavioral performance in visual acuity and face discrimination tasks, as well as increased structural integrity in late-visual pathways, partially moderating the behavioral improvement (Pedersini et al., 2023). Together, these findings suggest that gray matter changes are responsive to congenital visual deprivation, with alterations extending beyond the early visual cortex to include parietal and frontal regions, rather than directly reflecting the effects of sight-restorative eye surgery. In contrast, white matter tracts appear to serve as more sensitive markers of brain plasticity following vision restoration.

## Discussion

Early blindness significantly reshapes the brain’s structural and functional organization. While most studies have focused on cortical thickness and volume, few have investigated changes in cortical myelin content. Our study expands upon these traditional approaches by examining both gray matter volume and myelin content in the same clinical population. A critical question remains as to whether these brain alterations can be reversed after sight-restorative surgery. Here, we investigated anatomical changes in individuals who have been treated for dense bilateral congenital cataract, and who showed increased integrity in late-visual white matter pathways (Pedersini et al., 2023).

### Assessment of structural alterations following congenital blindness

In our study we observed structural plasticity following congenital visual deprivation, mainly affecting cortical thickness and T1-w signal.

In terms of cortical thickness, we found thicker cortex primarily in posterior brain regions, likely due to reduced synaptic pruning of exuberant connections (such as axon collaterals and dendritic arbors) (Jiang et al., 2009) or an increase in glial proliferation following sensory deprivation during the sensitive period of visual development. This finding contrasts with the cortical thinning of occipital regions (V1 to V5) typically observed in amblyopia (Liang et al., 2019). Our ROI-level analysis revealed two regions with thicker cortex (CT) in CC individuals: the lingual gyrus and the rostral anterior cingulate cortex (rACC), aligning with previous findings (Anurova et al., 2015; Inuggi et al., 2020). Interestingly, (Du et al., 2009) reported cortical thinning in the lingual gyrus in amblyopic patients, which contrasts with our results. These discrepancies may reflect differences in underlying developmental mechanisms. In congenital bilateral cataracts, the complete visual deprivation during sensitive periods leads to a lack of sensory-driven synaptic pruning in the occipital cortex, resulting in a thicker cortex. In amblyopia, however, partial visual input from the weaker eye triggers selective suppression by the brain, leading to disuse-driven pruning and reduced cortical thickness, to optimize the processing of the stronger eye’s input. In addition, we observed greater CT in normal sighted controls (NSC) within the temporal pole, a complex anatomical area involved in high-level cognitive and visual functions, such as visual scene analysis, face recognition, and visual memory (Herlin et al., 2021).

It is essential to consider that studies in healthy populations have consistently reported a negative correlation between cortical thickness and BOLD response magnitude, indicating that a thicker cortex is generally less strongly activated than a thinner one (Lu et al., 2009; Nuñez et al., 2011). This relationship may be explained by the synaptic pruning process, which strengthens activation through selective synaptic refinement. In blind individuals, previous findings from Anurova et al. (2015) suggest a similar negative correlation between CT and BOLD response amplitude in occipital regions during auditory tasks, that mirrors the pattern seen in NSC across task-related areas. This implies that the occipital cortex may play a functional role in higher-order processing of auditory information. Although task-related data were unavailable for this clinical group, we hypothesize that the thicker cortex observed in occipital regions might correspond to reduced functional activation in these areas during non-visual tasks.

With respect to cortical myelin content (approximated by T1-weighted signal), we observed greater T1-w signal in multiple regions across occipital, frontal, and parietal lobes. This distribution partially overlaps with regions showing an increased degree of cortical folding, revealing a more extensive pattern of structural changes than previously reported (Voss et al., 2014). Specifically, our findings include a considerable number of regions beyond the occipital lobe, suggesting that sensory deprivation induces increased cortical myelin content across a broader network. In a recent study (Pedersini et al., 2023), we observed similar integrity in late-visual white matter pathways when comparing the same clinical population with controls, despite the lack of visual experience. Interestingly, several regions showing higher cortical myelin content are connected by late-visual pathways, particularly the Inferior Frontal Occipital Fasciculus (IFOF) (Conner et al., 2018) and Inferior Longitudinal Fasciculus (ILF) (Herbet et al., 2018), supporting effective communication within and between these cortical areas even under congenital sensory deprivation. This increased communication efficiency may represent the structural basis for the heightened functional interaction observed between occipital regions and higher-cognitive frontoparietal networks, known to support language, numerical cognition, executive control, and working memory (Bedny, 2017; Bedny et al., 2011; Deen et al., 2015; Pedersini et al., 2024).

### Assessment of the impact of sight-recovered surgery on brain structure

In the current study we do not only test for structural differences following the sensory deprivation (a *congenital deficit*), but we also track structural changes following sight-recovery in CC adolescents, that represents a unique case of a *benefit, acquired* later in life.

In the literature, several studies assessed functional plasticity following an *acquired deficit*, in multiple sensory modalities. In 2009, Wandell and Smirnakis report a lack of plasticity, in favor of stability, in the primary visual cortex, following retinal damage, either monocular or binocular. This finding has been confirmed in humans (Baseler et al., 2011; Huber et al., 2015; Levine & McAnany, 2005) and in animals (Smirnakis et al., 2005) and align with the idea that longer-term structural plasticity is negligible in early visual cortex and increases along the visual pathways (Haak & Beckmann, 2019). Interestingly, this lack of functional plasticity following an *acquired deficit* might be a specific feature of the visual system (late blind individuals), as some studies (Buonomano & Merzenich, 1998; Nahum et al., 2013) reported functional plasticity in the motor cortex in animals even following an *acquired deficit* (limb amputation).

Our results build upon these findings by exploring the structural implications of a benefit acquired during adolescence (i.e., *sight recovery*) rather than a deficit (i.e., *sight loss*). We demonstrate that the structural changes resulting from congenital sensory deprivation do not revert after eye surgery performed once the sensitive period has closed. These findings highlight that, irrespective of the underlying brain mechanisms - whether synaptic pruning or excessive glial proliferation - none of them significantly revert following the sight-recovery surgery. Interestingly, in the same clinical population, we observed brain plasticity in cortico-cortical late-visual white matter, partially moderating the increased behavioral performance, even though the eye surgery was conducted during adolescence (Pedersini et al., 2023). This may indicate that white matter serves as a more reliable biomarker of brain plasticity in cases of *benefits*, such as *sight recovery*, acquired during adolescence.

#### Limitations

It is important to mention two main limitations of this study. First, despite the substantial number of patients in our clinical sample and the longitudinal nature of the study, it is important to notice that the number as well as the time window between structural data acquisitions differs among patients, due to the challenging conditions of data collection.

A second important limitation of this study is linked to the inclusion of controls and is represented by the socio-economic differences between CC and NSC. While it is essential to account for these differences, we have taken steps to control for these by normalizing structural measures using intra-subject z-scores. The consistency of our results with previous research supports the robustness of our approach and reinforces confidence in the validity of our findings.

## Acknowledgements

NYUAD Center for Brain and Health, funded by Tamkeen under NYU Abu Dhabi Research Institute grant CG012. The ASPIRE Precision Medicine Research Institute Abu Dhabi (ASPIREPMRIAD) grant number VRI-20-10, funded by ASPIRE, the technology program management pillar of Abu Dhabi’s Advanced Technology Research Council (ATRC).

