## Supplementary Material for "Gray matter abnormalities in sight deprivation and sight restoration"

**Affiliations:**


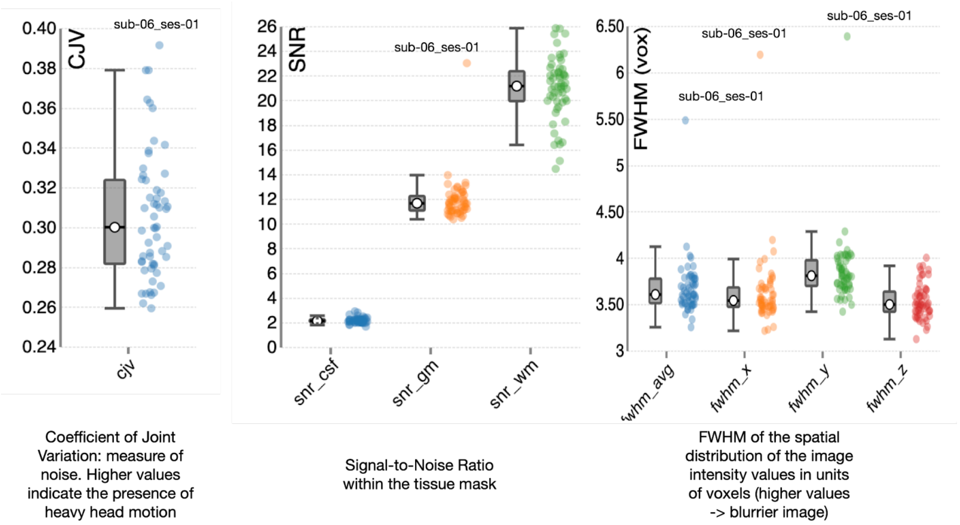


**Fig. S1.** Section of the report of structural images quality from MRIQC.

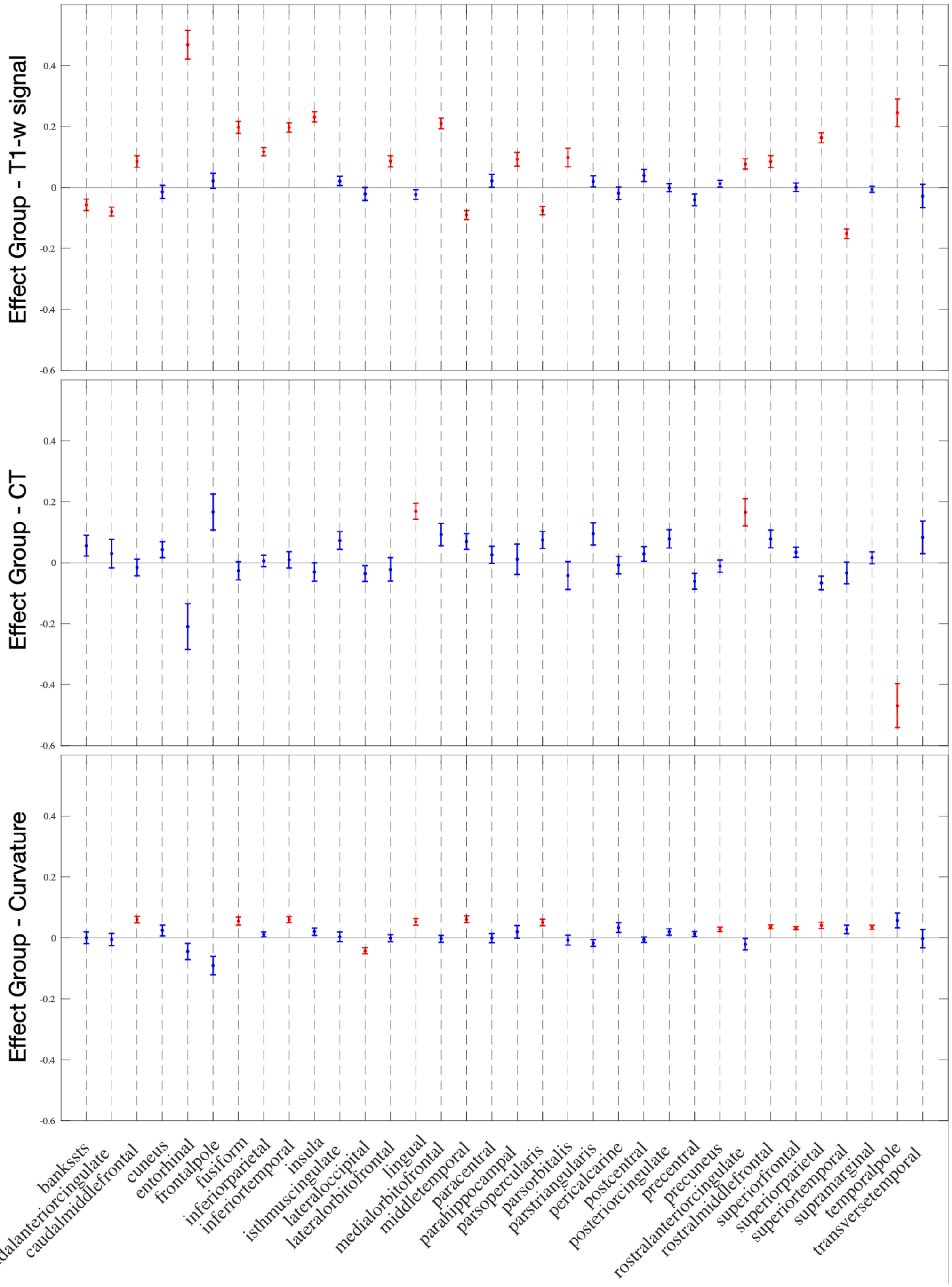


**Fig. S2. Effect of group at ROI-level.** Betas representing the group effect for T1-w signal intensity, cortical thickness, and curvature across different ROIs. Red bars indicate significant results, while blue bars denote non-significant findings. Error bars represent the standard error around each beta estimate.

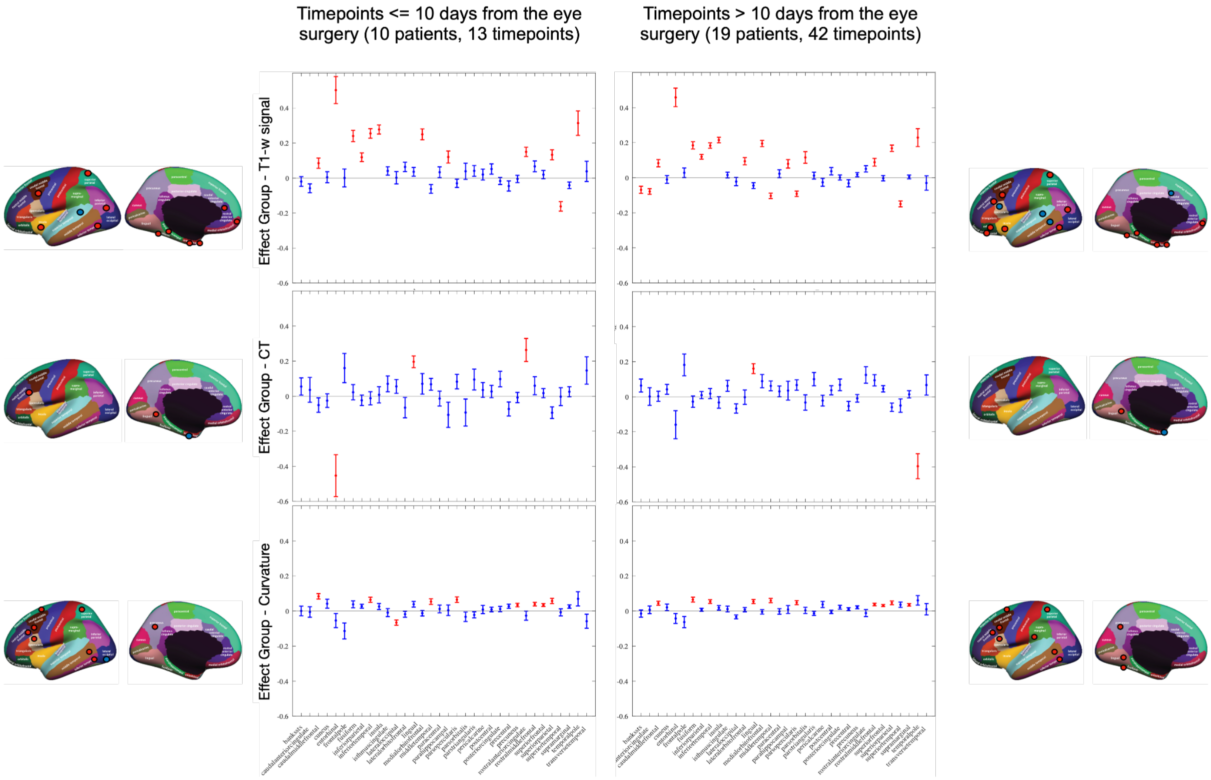


**Fig. S3. Effect of group at ROI-level including only timepoints around the surgery (left column) or significantly after the surgery (right column).** Central columns – Betas representing the group effect for T1-weighted signal intensity, cortical thickness, and curvature across different ROIs. Red bars indicate significant results, while blue bars denote non-significant findings. Error bars represent the standard error around each beta estimate. Lateral columns – Visualization of the Desikan-Killiany ROIs on medial and lateral brain surfaces. Red dots highlight regions with significantly higher betas in CC individuals, whereas blue dots indicate regions with significantly higher betas in NSC.

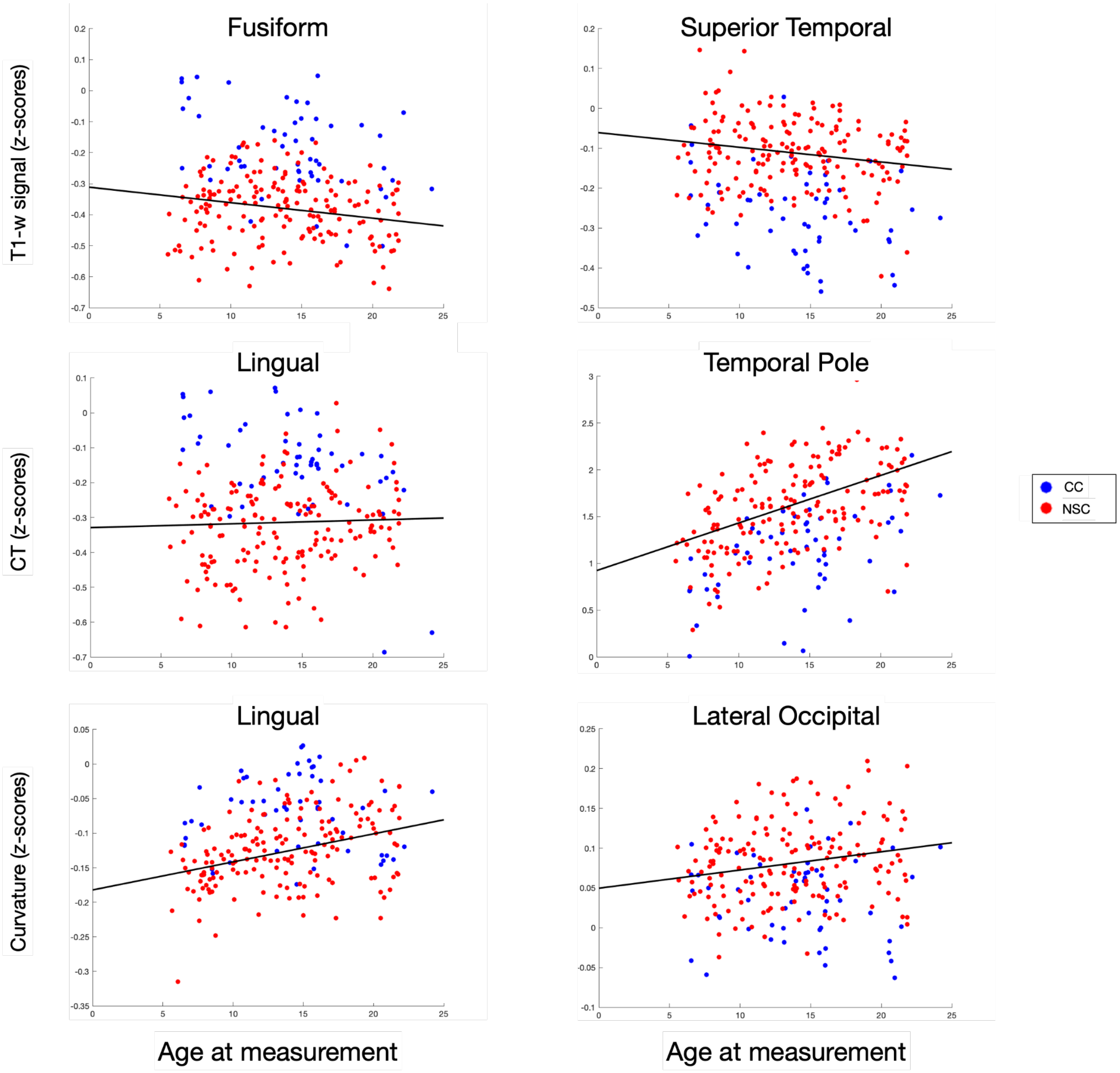


**Fig. S4. T1-w signal, CT and Curvature as a function of age.** For each structural metric, we selected two representative regions of interest (ROIs) and represent the trend of the structural metric as a function of age, for each group (CC in blue and NSC in red). Left panel: Regions with higher z-scores in congenital cataract-reversal (CC) individuals compared to normally sighted controls (NSC). Right panel: Regions with higher z-scores in NSC compared to CC. The black line in each plot represents the age-related effect (beta coefficient) on each measure.

In Fig. S5, we illustrate the pattern of changes in T1-w signal as a function of age within the regions of interest (ROI) where we observed a significant effect (FDR-$p < 0.01$). Research indicates that absolute T1-w signal trends typically follow an inverted U-shaped pattern: younger individuals (approximately 20 years old) exhibit lower T1-w signals, which stabilize around ages 40 to 50 before beginning to decline again (Rowley et al., 2017). However, data on individuals aged 0 to 20 years are limited. Importantly, we are utilizing z-scores, which emphasize effects relative to the mean rather than absolute changes. This approach allows us to identify ROI-specific effects that may not be apparent when examining general trends of absolute values. Regarding our results, the dorsal locations align with the overall observation of an increasing T1-w signal with age. Since our sample does not extend into the stabilization phase (40-50 years), we only observe the initial increase. In contrast, the ventral location displays a decreasing trend in T1-w signal with age. Given that we are analyzing relative signals, this decrease could indicate a relative decline within the context of an overall absolute increase in T1-w signals, or it may signify a decrease in absolute terms altogether. In conclusion, our findings represent an initial step toward bridging the knowledge gap regarding T1-w signals in individuals aged 0 to 20 years.

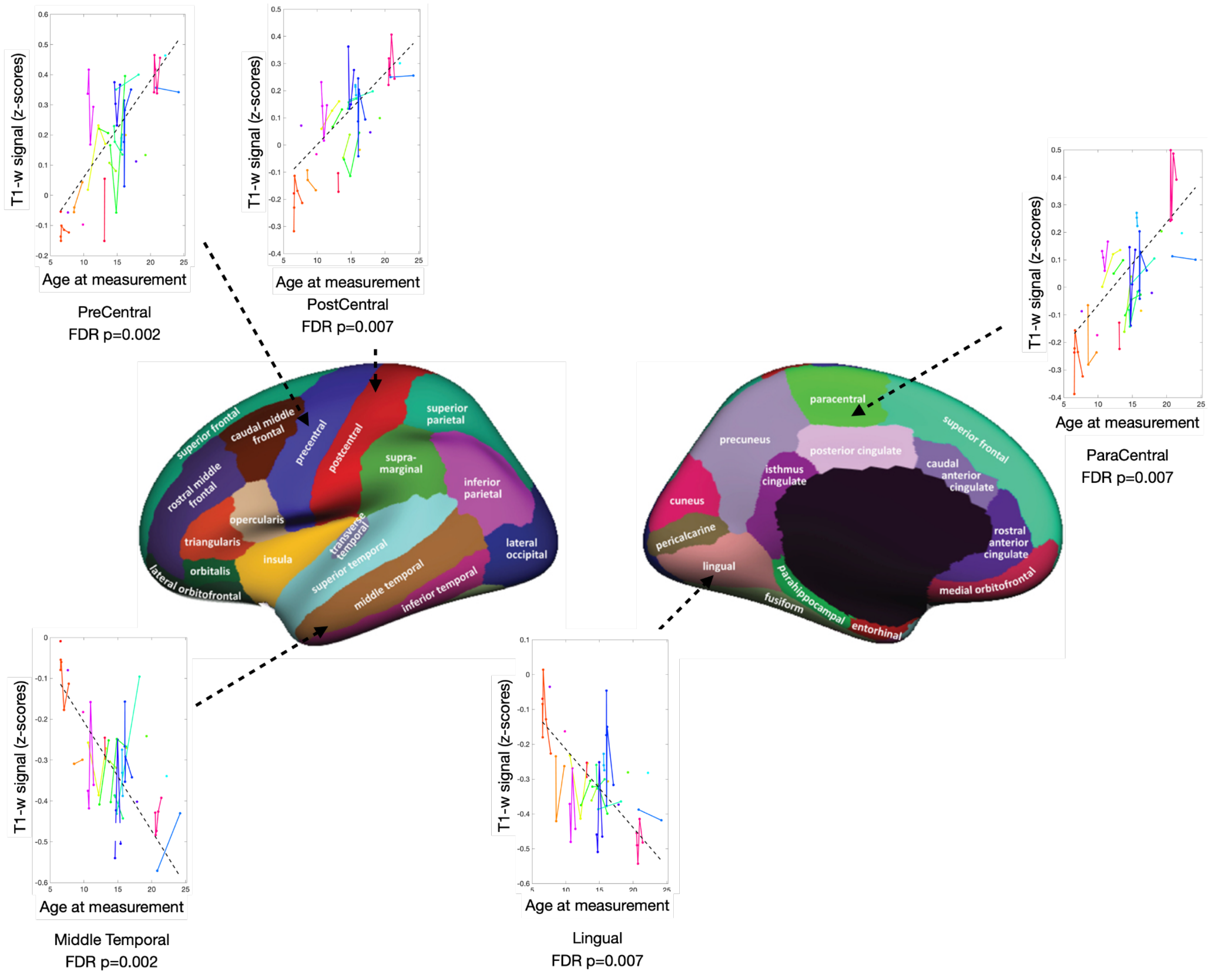


**Fig. S5. Effect of age at ROI-level in the group of individuals who have been treated for dense bilateral congenital cataract.** Lateral and medial brain surface superimposed with the Desikan-Killiany Atlas. Each plot represents z-scores T1-w signal for each subject as a function of age. Each color represents a CC and each line is connecting the multiple timepoints from the same subject. The dashed line indicates the beta extracted from the linear mixed-effect model, with the random-intercept of subjects. The plots have been chosen based on the ROIs reporting significant results after FDR.
